# Intra-host SARS-CoV-2 evolution in the gut of mucosally-infected *Chlorocebus aethiops* (African green monkeys)

**DOI:** 10.1101/2021.08.22.457295

**Authors:** Lori A. Rowe, Brandon J. Beddingfield, Kelly Goff, Stephanie Z. Killeen, Nicole R. Chirichella, Alexandra Melton, Chad J. Roy, Nicholas J. Maness

## Abstract

In recent months, several SARS-CoV-2 variants have emerged that enhance transmissibility and escape host humoral immunity. Hence, the tracking of viral evolutionary trajectories is clearly of great importance. Little is known about SARS-CoV-2 evolution in nonhuman primate models used to test vaccines and therapies and to model human disease. Viral RNA was sequenced from rectal swabs from *Chlorocebus aethiops* (African green monkeys) after experimental respiratory SARS-CoV-2 infection. Two distinct patterns of viral evolution were identified that were shared between all collected samples. First, mutations in the furin cleavage site that were initially present in the virus as a consequence of VeroE6 cell culture adaptation were subsequently lost in virus recovered in rectal swabs, confirming the necessity of this motif for viral infection *in vivo*. Three amino acid changes were also identified; ORF 1a S2103F, and spike D215G and H655Y, that were detected in rectal swabs from all sampled animals. These findings are demonstrative of intra-host SARS-CoV-2 evolution unique to this nonhuman primate species and may identify a host-adapted variant of SARS-CoV-2 that would be useful in future development of primate disease models.

## Introduction

The COVID-19 pandemic, caused by the coronavirus SARS-CoV-2, has killed more than 4 million people to date. Despite the development and rollout of safe and highly effective vaccines on an unparalleled time scale, nearly unchecked viral spread has continued due to vaccine refusal and hesitancy in nations with adequate vaccine supply coupled with inadequate supply in many regions of the world. As a result, several new variants have emerged with enhanced replicative or infectious capacity and immune escape [1]. Since the discovery of the D614G mutation [2,3], noted early in the pandemic and found to enable enhanced infection in cells and is now present in all sequenced isolates, several variants of interest and concern (as defined by the CDC) have arisen [1,4–11]. These include B.1.1.7 (alpha), which was originally detected in the United Kingdom and rapidly spread globally, B.1.351 (beta), originally detected in South Africa, and P.1 (gamma), originally detected in Japan in a traveler from Brazil. Most recently, the B.1.617.2 (delta) variant has rapidly become globally dominant, which is of great concern due to its greatly enhanced transmissibility relative to other variants and its ability to infect vaccinated individuals [12]. Collectively, these data suggest a concerning scenario wherein there is an ample ontological niche for continued SARS-CoV-2 evolution to facilitate persistence and pandemic spread in populations, including those with preexisting immunity to the virus.

Several animal species have been tested in efforts to develop a disease model that faithfully recapitulates human disease and associated pathological consequence, ultimately to be used in the evaluation of promising vaccines and therapies for COVID-19. Multiple nonhuman primate (NHP) species have been explored to this end, including *Macaca mulatta* (rhesus macaque) (RhM), *Papio anubis* (baboons) [13], *Macaca nemestrina* (pigtail macaques) (manuscript in preparation), *Macaca fascicularis* (cynomolgus macaque), and *Chlorocebus aethiops* (African green monkeys; AGM) [14–16]. Although some data suggest enhanced disease in AGM relative to other species [17], SARS-CoV-2 replicates to high titer in all of these species. However, in most cases, viral replication in upper respiratory sites is restricted to the first several days after infection. Thus, detection of intrahost evolution in these sites may be limited. Importantly, it is now becoming clear that the virus can persist and continue to replicate in gastrointestinal sites [18,19], which provides an opportunity to examine viral evolution beyond the first several days of infection and may illuminate tissue-specific compartmentalized evolution.

In this study, a focused set of gastrointestinal samples was used to examine viral and host dynamics in SARS-CoV-2 infected African green monkeys (AGM). Consistent patterns of evolution were identified, which is suggestive of species-specific adaptation. The samples were exclusively rectal swabs, which suggest viral replication in the gastrointestinal system may be an important source of viral variants. Notably, these results have important implications for the detection of SARS-CoV-2 variants in wastewater [20,21].

## Materials and Methods

### Sample collection, RNA Isolation and conversion to cDNA

Rectal Swabs were collected and stored in RNA/DNA Shield (Zymo Reseach, Irvine, CA). RNA was isolated using the Zymo Quick-RNA Viral kit and converted to cDNA using Protoscript II (New England Biolabs, Ipswich, MA) as follows: 10μl Template RNA, 1μl 10μM random hexamers, and 1μl 10mM dNTPs were incubated at 65°C for 5 minutes and then place directly on ice for 1 minute. The following was then added: 4μl PSII buffer, 2μl 100mM DTT, 1μl RNase inhibitor, and 1μl PSII reverse transcriptase and incubated at 42°C for 50 minutes, then 70°C for 10 minutes, followed by a hold at 4°C.

### Sequencing

DNA libraries were made using the standard SWIFT Normalase Amplicon Panels protocol (SWIFT Biosciences, Ann Arbor, MI) utilizing the SNAP UD indexing primers. The libraries were normalized to 4nM and pooled. Paired-end sequencing (2 × 150) was performed on the Illumina (San Diego, CA) MiSeq platform.

### Data analysis

Primer sequences were trimmed, and sequence reads were aligned to the SARS-CoV-2 genome (WA1/2020 isolate, accession MN985325) using the built-in mapping function in Geneious Prime software. Variants were called that were present at greater than 10% of reads at that site. Variants detected in the SARS-CoV-2 spike structure were visualized with UCSF Chimera, developed by the Resources for Biocomputing, Visualization, and Informatics at the University of California, San Francisco [22].

## Results and Discussion

A high-density overlapping amplicon approach was used to amplify and sequence the entire SARS-CoV-2 genome from the viral stock used to experimentally infect the AGM. Rectal swabs isolated from three AGM infection at 3-4 weeks post infection were interrogated in the same fashion. The challenge stock harbored several mutations of various frequencies relative to the WA1/2020 patient isolate (Figure 1A). These included two mutations in spike, F79L and a high frequency mutation that changed the arginine (R) at position 682 to a leucine (R682L) (Figure 1A). This residue is the second arginine, and one of the key residues, in the RRAR furin cleavage site, most likely the result of adaptation to VeroE6 cells. This stock was used to infect three AGMs from which samples were available. Samples were acquired from nasal, oropharyngeal, and rectal swabs from time points spanning four weeks postinfection. RNA extracted from swabs at early time points were exhausted from extensive qRT PCR characterization and focus was shifted to rectal swabs collected from late time points including 21 and 28 days post infection in three of the AGMs. Sequencing of these samples revealed a consistent pattern of evolution *in vivo* with three mutations arising in all three animals; ORF1a/b S2103F, spike D215G and spike H655Y (Figure 1B). In contrast, F79L and R682L mutations, present at approximately 60 and 80% in the virus stock, respectively, were completely absent in all rectal samples.

**Figure 1.**
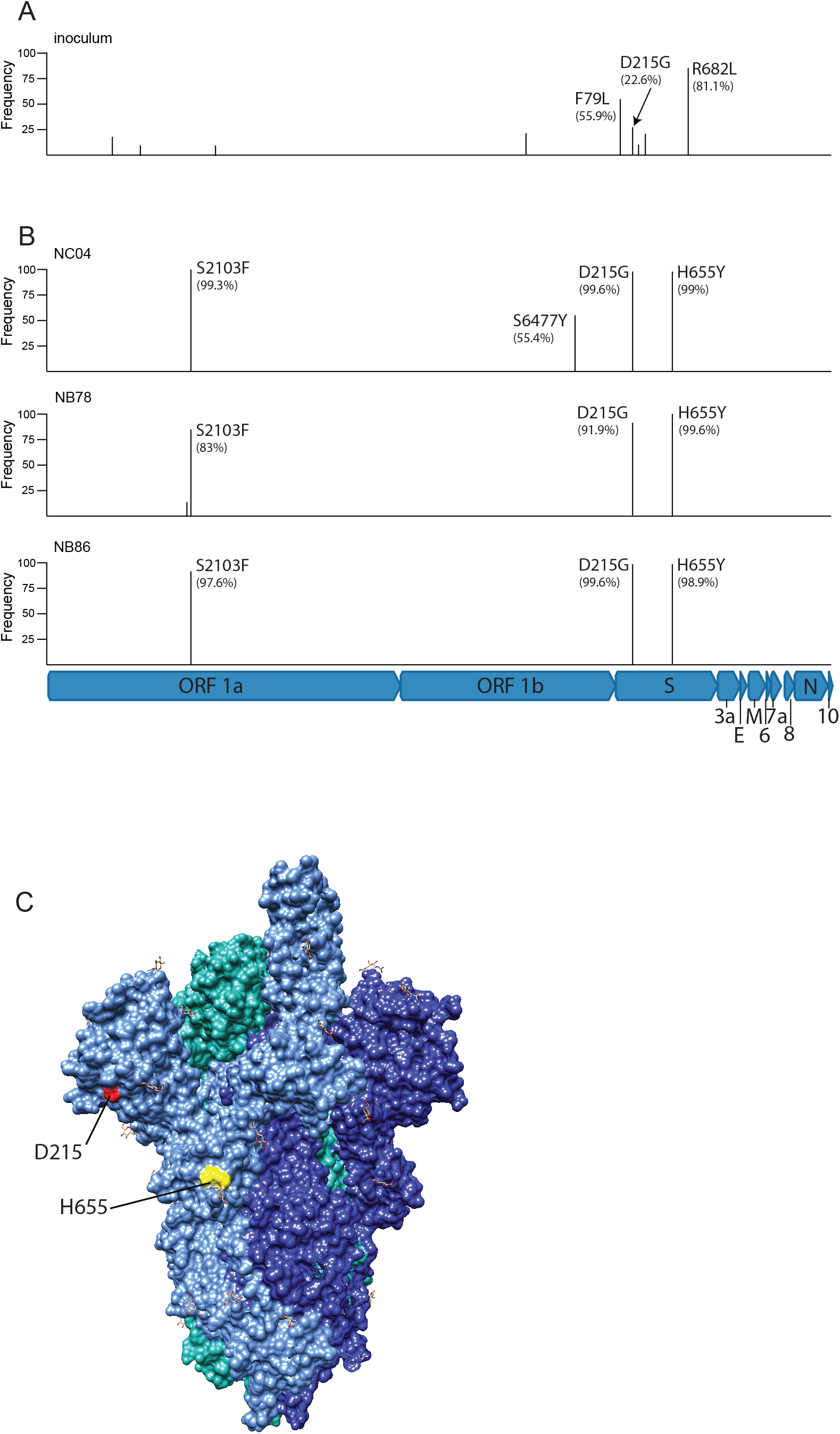
Viral evolution detected in rectal swabs in SARS-CoV-2 infected AGM. AGM were challenged via intranasal and intratracheal route with SARS-CoV-2 using a stock of the WA1/2020 isolate harboring several variants relative to the patient sample (A). Sequencing of viral RNA isolated from rectal swabs at 3-4 weeks after challenge revealed a consistent pattern of variants in all three animals (B). Two spike mutations detected in all animals map to the N-terminal domain (D215G) and near the furin cleavage site (H655Y) based on a spike structure (PBD 7K8Z [34]) (C).

The ORF1a/b residue 2103 lies in nonstructural protein 3 (nsp3), which is the largest protein in the SARS-CoV-2 proteome and localizes with other viral proteins to the cell membrane [23,24]. The S2103F mutation in this protein has been detected at low frequency in human samples including in India [25], but is not restricted to a particular clade and through the course of the pandemic has never been detected in greater than 1% of all sequenced isolates (nextstrain.org [26,27]).

The R682L mutation likely eliminates the furin cleavage site in spike. Reversion to the wild type amino acid, therefore, confirms the necessity of the furin cleavage site for infection *in vivo*, which has been previously suggested [28]. This finding, although not unexpected but noteworthy, confirms that the loss of the furin cleavage site *in vitro* can be ascribed to replication in VeroE6 cells [29]. Since VeroE6 cells are sourced from AGM kidney, this finding confirms the loss of the furin cleavage site to be an *in vitro* phenomenon and not specific to replication in AGM cells *per se*. The D215G mutation rose from approximately 15% prevalence in the stock to near 100% in all three animals. This mutation lies in the N-terminal domain of the spike protein (Figure 1C) and is detected at low frequency in multiple viral lineages and is a defining variant of the B.1.351 (beta) lineage (nextstrain.org). One report showed this mutation has a weak but detectable impact on augmenting cell to cell fusion [30]. The H655Y mutation which lies immediately adjacent to the furin cleavage site (Figure 1C) and has been detected sporadically in multiple viral lineages with the P.1 (gamma) lineage being most prominent as H655Y is a defining variant of the gamma lineage (nextstrain.org). This variant has also been detected in other animal species disease models including cats and mink [31,32]. A recent report also showed that virus harboring a 655Y amino acid showed enhanced replicative capacity *in vitro* and spike protein cleavage [32], likely explaining the H655Ymutation in the AGM species.

In this study, we define a consistent pattern of viral evolution in SARS-CoV-2-infected AGM. Despite a limited sample size, the ubiquity with which this pattern was detected strongly suggests a selective advantage to the identified mutations. All samples used in this study were derived from rectal rather than respiratory samples, and the data do not clearly suggest a possible mechanism for this selection. It is plausible that replication in the GI tract presents a unique set of selective forces on the virus driving evolution to the mutations identified. Long-term persistence of the virus in the gastrointestinal tract, as has been suggested for SARS-CoV-2 in humans and is a well-described phenomenon in other coronaviruses [33] may also allow selected mutations to accumulate. Even in the latter case, these data suggest the virus continues to replicate and evolve throughout the gastrointestinal system, which may further reveal novel aspects of the virus-host relationship and may have important implications for the detection of variants in wastewater.

## Conclusions

These data and study support two important conclusions. First, previous reports that an intact furin cleavage site is important for infection *in vivo* were confirmed in this study. Second, the combination of the three mutations identified (ORF1a/b S2103F, spike D215G and spike H655Y) represents a new host-adapted SARS-CoV-2 strain exclusive to the AGM species. Future experiments may include infection of AGM (or other nonhuman primate species) with this virus to optimize infectious challenge in product evaluation or immunopathogenesis studies. Alternatively, these mutations could be added to the backbone of variants of concern to assess their impact on experimental infection in AGM or other closely related nonhuman primate species.

## Author contributions

LAR, BJB, CJR, and NJM conceived and designed the experiments. LAR, BJB, KG, SZK, NRC, and AM performed the experiments and assisted with sample manipulation. NJM and CJR drafted the manuscript. All authors have read and agreed to the current version of the manuscript.

## Institutional Animal Care and Use Statement

The use of nonhuman primates in this project was reviewed and approved by the Tulane Institutional Animal Care and Use Committee (IACUC), protocol P0447.

